# Genomic from *Plasmodium falciparum* field isolates from Benin allows the identification of PfEMP1 variants by LC-MS/MS at the patient’s level

**DOI:** 10.1101/391615

**Authors:** Claire Kamaliddin, David Rombaut, Emilie Guillochon, Jade Royo, Sem Ezinmegnon, Stéphanie Huguet, Sayeh Guemouri, Céline Peirera, Cédric Broussard, Jules M Alao, Agnès Aubouy, François Guillonneau, Philippe Deloron, Gwladys Bertin

## Abstract

PfEMP1 are the major protein family from parasitic origin involves in the pathophysiology of severe malaria, and PfEMP1 domain subtypes are associated with the infection outcome. In addition, PfEMP1 variability in endless and current protein repository do not reflect the immense diversity of the sequences of PfEMP1 proteins. The aim of our study was to identify the different PfEMP1 variants expressed within a patient sample by mass spectrometry. We performed a proteogenomic approach to decipher at the patient’s level PfEMP1 expression in different clinical settings: cerebral malaria, severe anemia and uncomplicated malaria. The combination of whole genome sequencing approach, RNAsequencing, and mass spectrometry proteomic analysis allowed to attribute PfEMP1 sequences to each sample and classify the relative expression level of PfEMP1 proteins within each sample. We predicted PfEMP1 structures using the newly identified protein sequences. We confirmed the involvement of DBLβ in malaria pathogenesis and observed that CIDRα domains linked to ICAM-1 binding DBLβ domains displayed EPCR binding structure.

## Introduction

*Plasmodium falciparum* is one of the causal agents of malaria in humans and leads to the severe form of the disease. In endemic areas, children under 5 years old are victims of cerebral malaria (CM) and severe anemia (SA).Despite the introduction of intra venous artesunate therapy, severe malaria mortality remains significant −0.5% of malaria outbreak according to World Health Organization (WHO) (1).

Through its development in human erythrocytes, *P. falciparum* grows inside the erythrocyte and the infected erythrocyte’s (iE) surface changes in shape and rigidity. Parasite proteins exported at the host cell surface mediate infected erythrocyte’s adhesion to the host’s endothelium. This phenomenon, called cytoadhesion, is the main virulence factor of the parasite and contributes to the pathophysiology of severe malaria. The sequestration of infected erythrocytes in the capillaries leads to hypoxia, occlusion and endothelial activation. In CM pathophysiology, the sequestration of iE in the brain capillaries is believed to trigger coma and brain swelling (2).

Among the proteins exported at the erythrocyte’s surface, three variant surface antigens (VSA) families have been described: the repetitive interspersed family (RIFIN) (3, 4), subtelomeric variable open reading frame (STEVOR) (5, 6) and *Plasmodium falciparum* Erythrocyte Membrane Protein 1 (PfEMP1) (7), which is the most described VSA family. They are encoded by the multigenic *var* gene family (8–10), consisting in ~ 60 copies per parasite genome. The diversity among *var* sequences is almost endless (11, 12) thus participating to the infected erythrocyte ability to evade the immune system.

PfEMP1 proteins are high molecular weight transmembrane proteins (200-350 kDa), and are composed of an intra-erythrocytic segment, which is conserved, and a highly variable extracellular segment (13). The extra-erythrocytic segment is composed of 4 to 9 alternated Duffy Binding Like (DBL) or Cystein Inter Domain Rich (CIDR) domains. The nature and the arrangement of these domains determine the binding phenotype of the infected erythrocyte (13, 14).

Among the PfEMP1 receptor in human endothelium, the most common is the broadly expressed in human cells CD36, but it is not related to any specific form of malaria. In the context of severe malaria, two human host receptors for PfEMP1 binding have been identified: the InterCellular Adhesion Molecule-1 receptor (ICAM-1) (15) and the Endothelial Protein C Receptor (EPCR) (16), both expressed by brain endothelial cells (17), and co-localized with the sequestered iEs (15). Specific domains from PfEMP1 are known to interact with ICAM-1, including the Domain Cassette 4 (DC4) (DBLα1.4/1.1-CIDRα1.6-DBLβ3) (18). The binding domain is located in the C-terminal third of the DBLβ3 (18). A study conducted on lab-adapted patient’s isolates after selecting on ICAM-1 highlighted that the residues involved in PfEMP1 binding to ICAM-1 are highly variable with a limited binding pattern (19) (20).

EPCR role in PfEMP1 binding has been more recently discovered (16). EPCR binding is mediated by highly variable but structurally conserved CIDRα1 PfEMP1 domains (more precisely CIDRαl.1 and CIDRal.4-1.8) (21, 22). Importantly, the level of PfEMP1transcript associated with EPCR binding is higher in samples from patients suffering from severe malaria, and increases with the disease severity (21).

A dual-binding with EPCR and ICAM-1 has been suggested, since not all CM isolates present an increase in binding-EPCR PfEMP1 coding transcript (23). In addition, expression levels of *var* coding transcripts are increased in parasites able to bind to EPCR, ICAM-1 and CD36 *in vitro* enforcing the idea of a mutual binding (24). The expression of DBL involved in ICAM-1 binding is associated with dual ICAM-1 and EPCR binding (20).

Despite their high diversity, PfEMP1 proteins are a target of naturally acquired immunity against *P. falciparum* (25), and the parasite express VSA corresponding to gap in the antibodies repertoire from an individual (26, 27). Therefore, identification and characterization of severe malaria associated PfEMP1 proteins is crucial. It is believed that despite the endless variability of PfEMP1 proteins (28), a cross-reactivity between expressed epitopes exists (29), as children in endemic area acquire protective immunity to the high binding phenotypes after a limited number of infection compared to PfEMP1 variability (26). PfEMP1 proteins are a proposed target for vaccine to protect people against severe forms of malaria (25, 30).

Most field studies looking for *P. falciparum* binding phenotypes are based on molecular biology analysis, and have shown that transcript coding for specific PfEMP1 domains expression level are associated with disease outcome (21, 24, 31, 32). However, this strategy is currently limited to the already identified PfEMP1 domains and does not give proficiency of the expressed proteins.

To complement this deficiency, mass-spectrometry-based proteomics is a powerful and sensitive tool for bottom-up protein identification after trypsin digestion. The application of mass spectrometry for VSA identification, and more especially PfEMP1 identification remains challenging for the following reasons. First, PfEMP1 are transmembrane proteins, and therefore present hydrophobic segments which decrease its solubility and its accessibility to proteases such as trypsin. Second, PfEMP1 are high molecular weight proteins (250-350 kDa), and finally, PfEMP1 are highly variable in sequences, yet database repository is usually simplified by eliminating redundancy. Thus, they do not reflect the natural sequence diversity which may occur in such a context.

To date, PfEMP1 identification with mass-spectrometry remains limited. In fact, the identification of the semi-conserved variant VAR2CSA in the context of Pregnancy Associated Malaria has been performed (33), but the identification of PfEMP1 variant in a context of severe malaria (CM or SA) has not been reported yet. The association of PfEMP1 domains with clinical outcome (UM, CM and Pregnancy Associated Malaria) was performed, with relaxed analysis parameters to allow the identification of PfEMP1 domains (34).

To identify PfEMP1 associated with *P. falciparum* clinical outcome in endemic settings, we used a “proteogenomic” strategy on field sample from children from Benin, West Africa, retrieved from May 2016 to August 2016. Specific PfEMP1 sequences from each isolate were reconstructed, using whole genome sequencing (WGS) data to enrich the protein database (Figure 1). We analyzed the whole proteome of samples from patients presenting either CM, SA or UM, and attribute PfEMP1 sequences within these samples. Corresponding samples were analyzed in RNAsequencing for PfEMP1 expression analysis, in relation to proteomic results.

**Figure 1:**
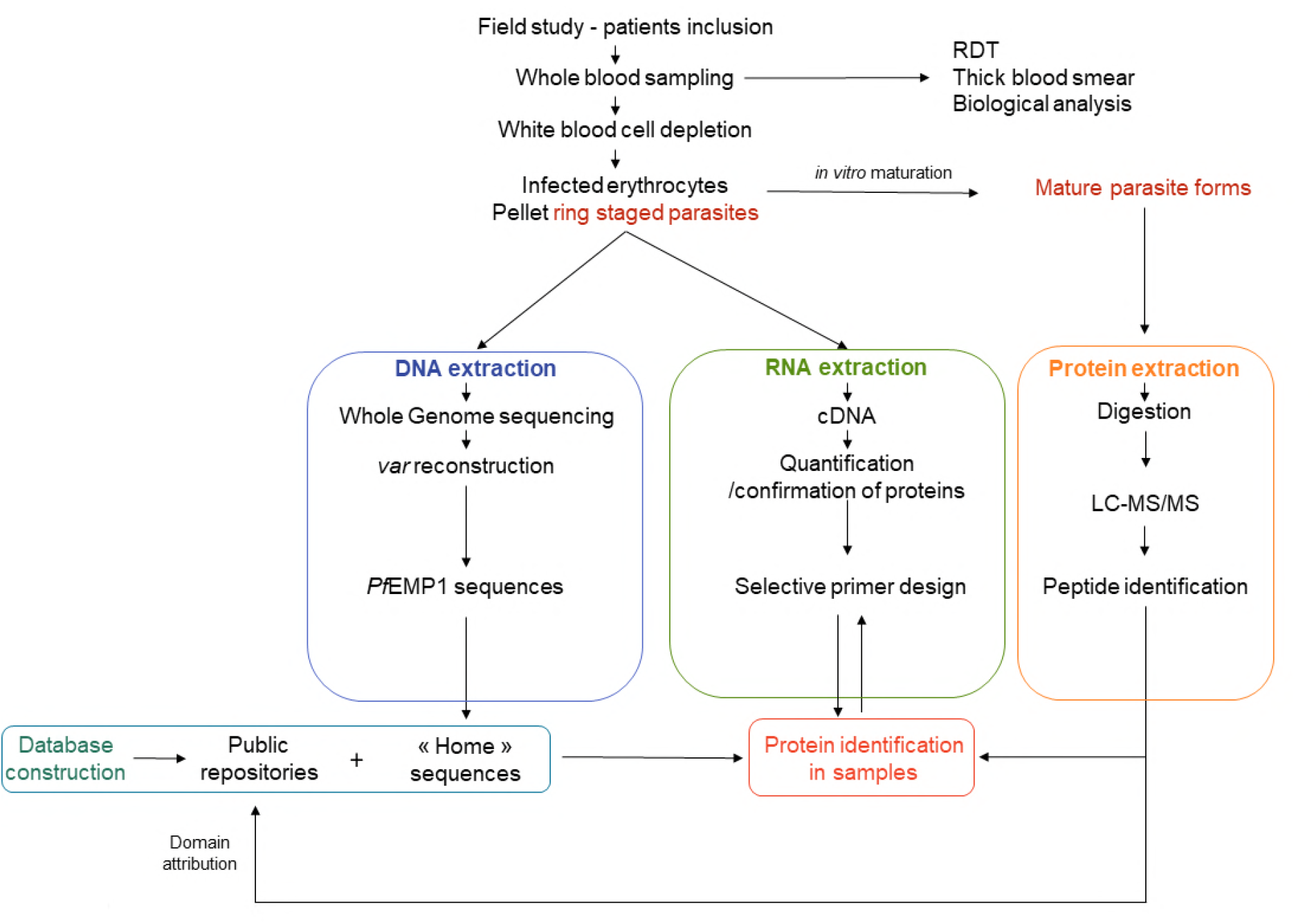
Proteogenomic approach on field samples for PfEMP1 identification. Whole blood sample from patients are collected. DNA and RNA are extracted from parasite’s ring forms. From the same collected tube, parasites are matured and the corresponding proteins from mature form (within one maturation cycle) are extracted and analyzed in LC-MS/MS. Whole genome sequencing data enrich the protein database used for protein identification with LC-MS/MS data. Selective mRNA sequencing provides additional information’s to elucidate proteins identification within the home-made database.

We managed to identify the PfEMP1 sequence associated with 4 CM samples, 9 SA and 9 UM samples using mass spectrometry (MS). For two samples, the relative expression levels from the PfEMP1 identified in MS was assessed using RT-qPCR using the corresponding mRNA. We confirmed the expression of several PfEMP1 within a single field isolates and provided the first identification at the patient’s level of PfEMP1 expressed by the parasite in the context of acute *P. falciparum* infection. We were also able to predict PfEMP1 structure within each newly identified sequence.

## Results

### Included Samples

We included 95 patients, covering 31 SA, 18 CM and 46 UM. Average patient’s age was similar among all inclusion groups. Parasite density geometric mean was 8,055 p/μL for UM group, 34,191 p/μL for CM and 24,313 p/μL for SA. Parasite density was only significantly different between SA and UM samples (*p* = 0.0158 with Bonferroni’s Multiple Comparison Test). Hemoglobin level was measured for 20 UM, 17 CM and 31 SA, respectively 11.28 [10.26; 12.75], 5.51 [4.10; 6.56] and 4.393 [3.90; 5.00] g/dL. Hemoglobin level was statistically different for UM samples (vs. CM and *vs*. SA) (*p* < 0.05), but no difference was observed between SA and CM samples. No difference in erythrocyte count (*p* = 0.1274) and temperature (*p* = 0.9125) was retrieved between SA and CM. All CM patients presented a coma (average BS 2 [2;2]), while SA patients did not (average BS 4.6 [4;5]) (*p* < 0.0001). 14/18 CM patients presented convulsions, and 7/38 SA patients (*p* < 0.0001).

For mass spectrometry analysis, we selected samples among those showing successful maturation. Four CM, 9 SA and 9 UM samples have been further investigated. RNAsequencing was successfully performed on 10 samples corresponding to the one studied in mass spectrometry (1 UM, 5 SA (3 patients) and 4 CM (2 patients)). (Supplementary Table 1)

### *Var* genes transcripts identification with RNAsequencing

We then focused on a set of 120 proteins from PlasmoDB reference, corresponding either to PfEMP1, or to other proteins associated with knobs formation (Supplementary data Table 3). The presence of skeleton binding protein 1 (Pf3D7_0501300) and EMP1 (Pf3D7_0730900) was assessed in all samples except SA09. Focusing on PfEMP1, we observed that the conserved ATS coding region was the only site for PfEMP1 mRNA identification using reference sequences from PlasmoDB. We then performed a selective mapping of RNAsequencing reads using the sequences from *var* genes reconstruction, which allowed to identify the expressed *var* transcript for 3 samples (Supplementary Table 4).

### Protein identification

Protein identification was performed using a home-made database (reference sequences from human and *P. falciparum* repositories, and the assembled *var* from field samples) containing 295,601 protein sequences, among which 87,489 were *P. falciparum-associated* sequences, while the other referred to the human proteome. Overall, we identified 3,300 proteins. After applying contaminant depletion and identification criteria, 3 214 proteins remained. A total of 1,302 were associated to the human proteome, and 1,912 to *P. falciparum*’s. Among those later, 460 proteins were identified as *P. falciparum* membrane-associated proteins, including 60.4% of hypothetical or putative, 12 % of PfEMP1s, 3.5% of RIFINs, 0.9% of STEVORs, 1.5% of PHISTs and 21.7% belong to other protein families.

A total of 155 proteins associated with PfEMP1 were identified, among which 96 were unique proteins, and clustered in 55 protein groups. Only 10 of the identified PfEMP1 (as part of protein group, or majority protein) were known sequences from public database repository (Uniprot and PlasmoDB). All other identified PfEMP1 sequences resulted from the translation of the reconstructed *var* genes from our samples.

### PfEMP1 identification and composition

We then focused on the 55 protein groups associated to PfEMP1 proteins. The minimum peptide number for protein identification was 2. Sequence coverage ranged from 0.6 to 24.6 % maximum. Average molecular weight of the identified PfEMP1 was 228.3 kDa. Using the VarDom online server, we reconstructed the domain architecture from the identified proteins (Figure 2). Fifty-three sequences displayed at least one hit with the repository sequences. NTS domain was found in 73.6 % of the sequences (39/53 sequences presenting at least one hit with VarDom reference sequences) and 97 % (38/39) of the sequences presenting NTS displayed the following combination: NTS-DBLα-CIDRα. The three-major head-terminal domain organizations were the following: NTS-DBLα-CIDRα-DBLβ (58.9%; n = 23/39), NTS-DBLα-CIDRα-DBLδ (30.8%; n = 12/39) and NTS-DBLα-CIDRα-DBLγ (2.6%; n = 1/39). To attribute more precisely subdomains to the identified PfEMP1 proteins, we performed a local blast from the 110 peptides associated to the PfEMP1 proteins against the VarDom sequences repository (Figure 3 and Figure 4). Fifty four percent (n = 60/110) of the peptides displayed at least one hit with the reference sequences from VarDom (median hit number per peptide was 5 (interquartile range [2; 23])), among which 40/110 peptides displayed hits with more than 90% identity with a reference sequence (among these, median number of hits per peptide was 2 (interquartile range [1; 5])). Most of the obtained hits from peptides with reference sequences corresponded to ATS sequences (171 hits), and 71 hits covering DBL domains (among them, 25 for DBLα, 20 for DBLβ, 14 for DBLδ, 1 for DBLε and 11 for DBLγ). One hit corresponded to NTS domain. Subdomain attribution was performed successfully for the corresponding peptides. No peptide was issued from CIDRα and seven peptides where issued from DBLβ.

**Figure 2:**
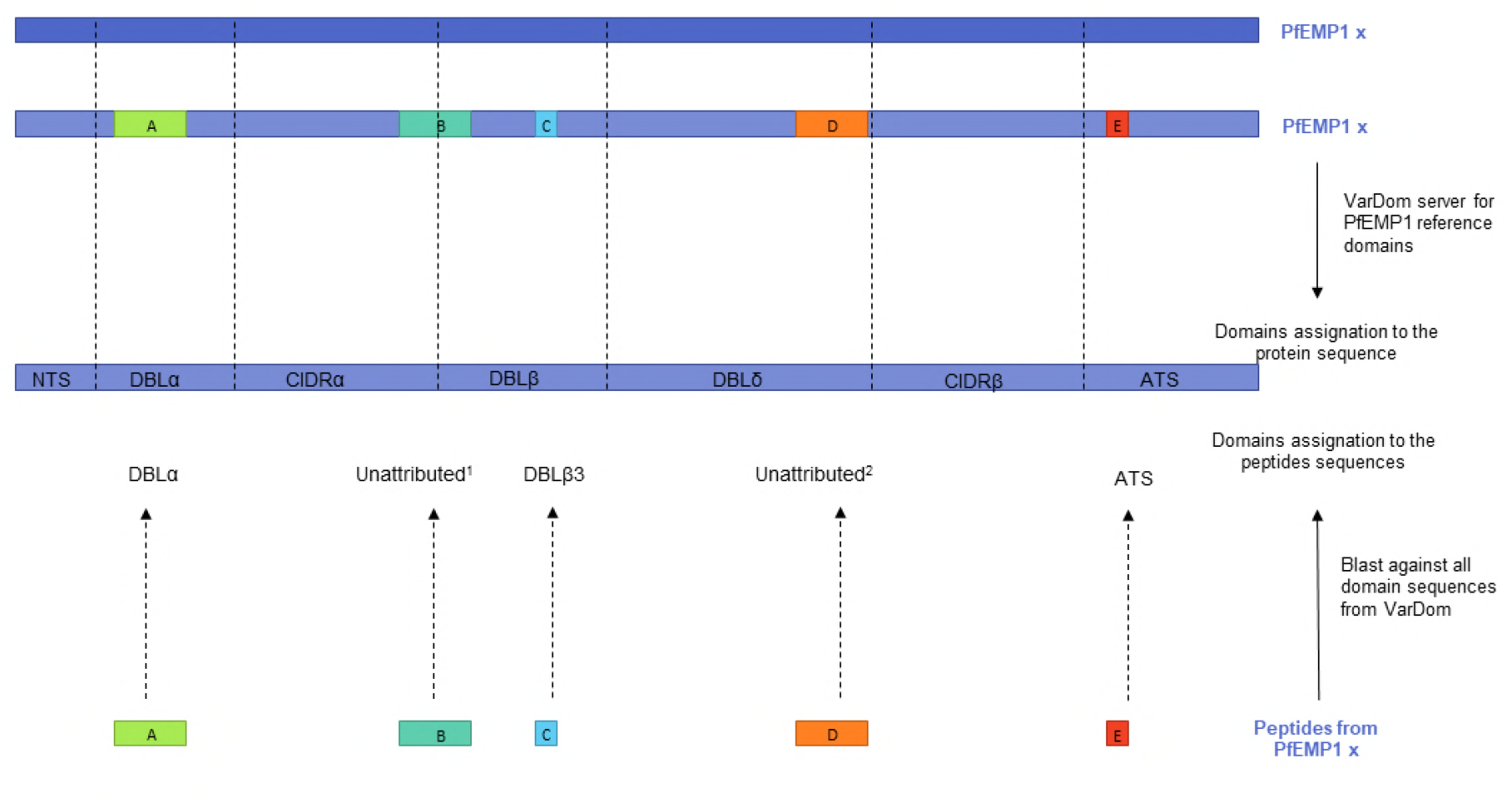
Schematic representation of domain attribution to PfEMP1 sequences and peptides. The PfEMP1 « x » has been identified in mass spectrometry with the peptides « A », « B », « C », « D », and « E ». The general domains organisation of PfEMP1 « x » can be attributed using the VarDom online servers. However, subdomains attribution is not over possible with confident score. Local blast of the peptides used for protein identification retrieves different types of results. First option (peptides « A », and « E ») is that the peptide local blast allows for a « general » domain attribution, without subtype. Second option (peptide « C ») is that the peptide is known within the precise sequence of a subdomain and allows identification of a subdomain. Unattributed peptides within the reference sequences has two origins. 1/The peptide is shared between two domains and the amino acids belonging to each domain are not sufficient for confident attribution. 2/The peptide is « unknown » in the reference sequence repository (a new PfEMP1 related peptide has been implemented).

**Figure 3:**
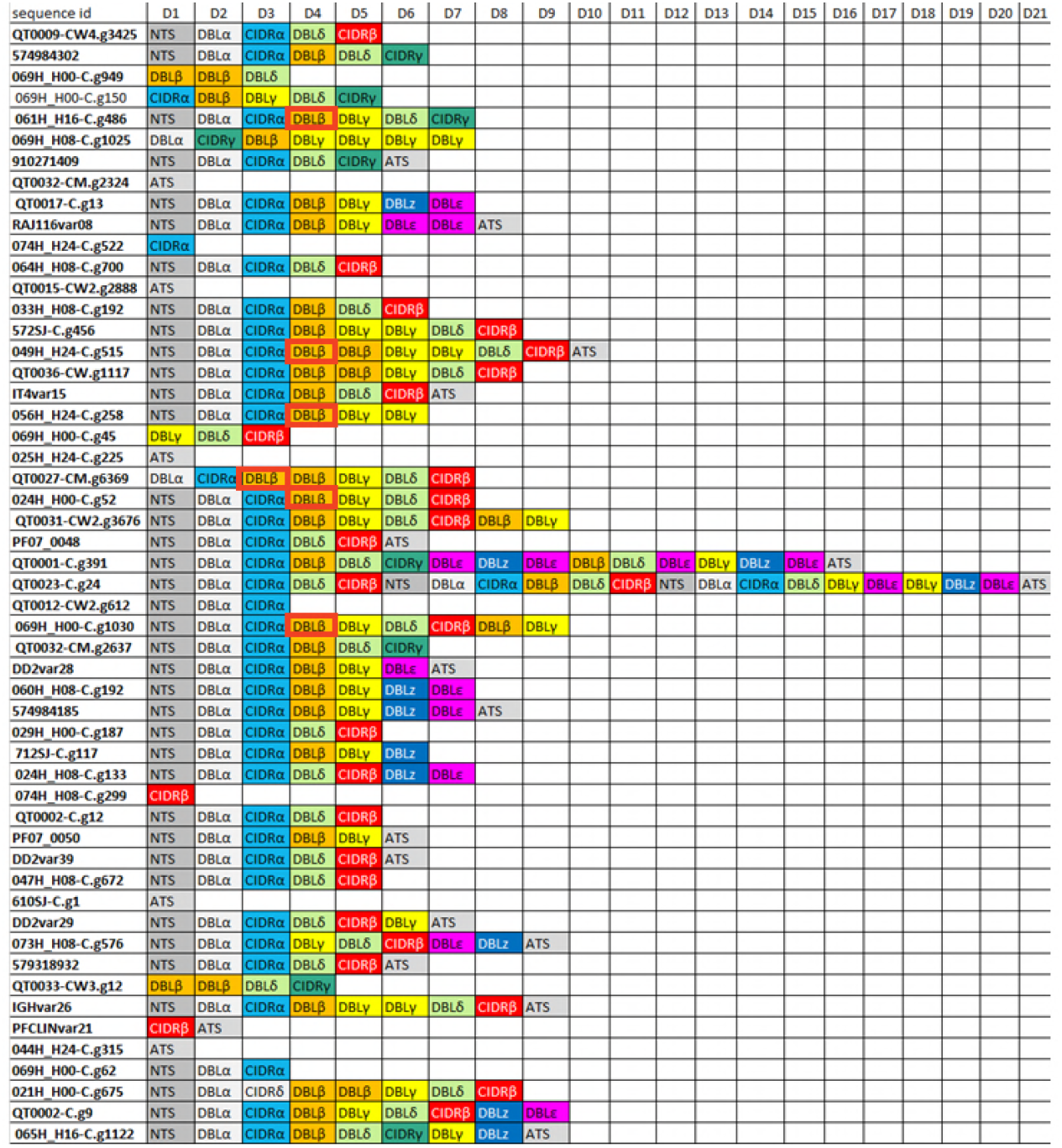
Domain description of the identified PfEMP1. PfEMP1 domain assignment for the one identified through mass spectrometry experiment. Proteins sequences have been mapped on VarDom server for domain attribution. Each color correspond to a domain type, corresponding to the VarDom server color code. DBLβ containing the binding pattern to ICAM-1 are highlighted with bold red rectangle.

**Figure 4:**
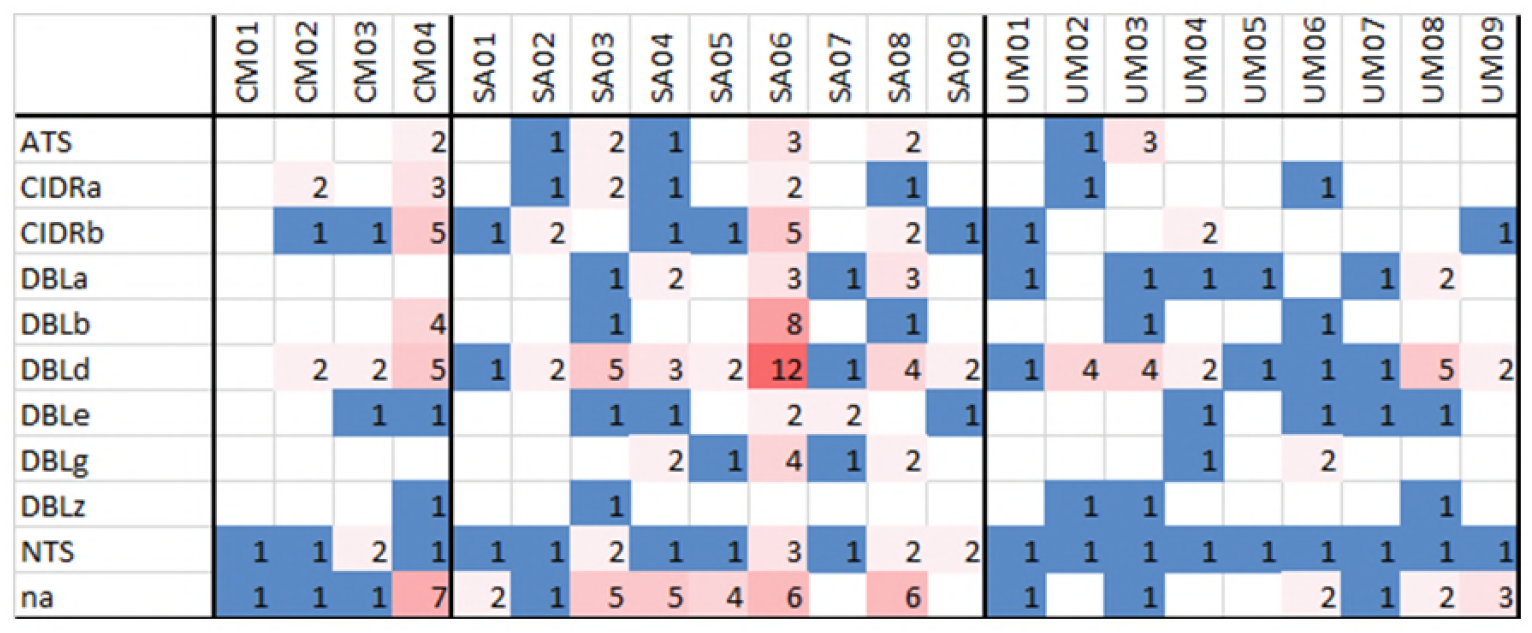
PfEMP1 domain count in the corresponding samples. Number of each domain attribute to the PfEMP1 peptides identified in CM, SA and UM samples. Color code is proportional to the number of identified domains (blue: low number and red: high number). NTS associated peptide is found once in each sample, except for 1 CM isolate and 4 SA isolates. CIDRα matching peptides have been observed in 2/4 CM, 5/9 SA and 2/9 UM. DBLδ matching domains have been found in 3/4 CM, 9/9 SA and 9/9 UM. In all samples, peptides corresponding to the protein sequence did not match with a known domain.

The specific search of the binding pattern for ICAM-1 retrieved 6 identifications within the PfEMP1 sequences identified in mass spectrometry.

### PfEMP1 associated with clinical outcome of infection

Among the identified PfEMP1 proteins in LC-MS/MS, 11 were shared between all patient’s group, 2 were specifically associated with CM, 5 with UM and 18 with SA. SA and CM shared 8 proteins, UM and CM 14 proteins, and SA and UM 6 proteins (Figure 5). The corresponding PfEMP1 are listed in Table 1.

**Figure 5:**
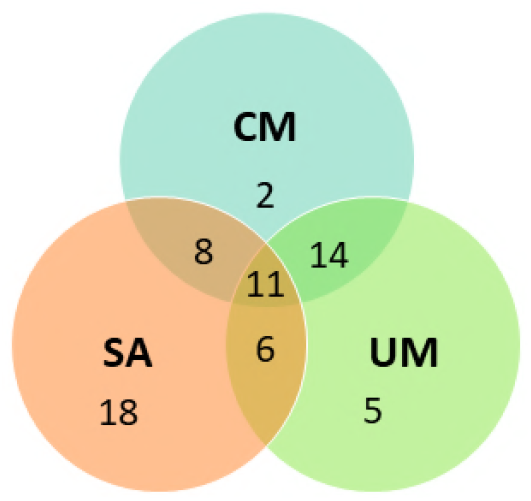
Venne Diagram representation of PfEMP1 identification. Representation of sequence identified in accordance with clinical presentation. 18 sequences were associated with SA only, 2 with CM and 5 with UM. SA and CM shared 8 sequences, CM and UM 14 sequences, and SA and UM 6 sequences. All three groups shared 11 sequences as well. CIDRα domain in found in ½ CM specific sequence; 13/18 SA sequences; 3/8 sequences shared between CM and SA and 3/5 UM sequences. Concerning DBLβ, the domain is contained in ½ CM sequences, 11/18 SA sequences, 4/8 sequences shared between CM and SA, and 4/5 UM sequences

**Table 1.**
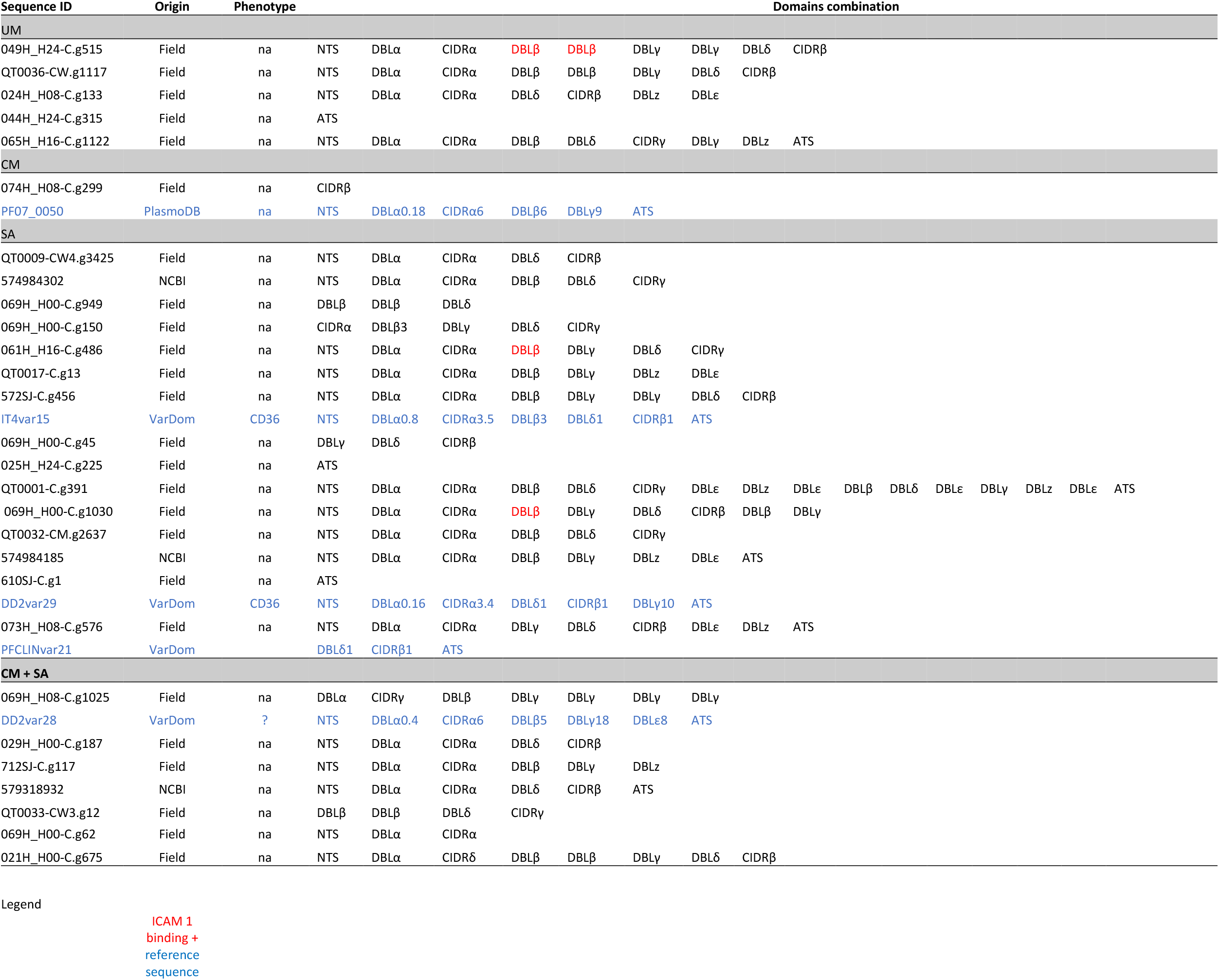
PfEMP1 sequences identified within each clinical group

| Sequence ID | Origin | Phenotype |  | Domains combination |  |  |  |  |  |  |  |  |  |  |  |  |  |  |
| --- | --- | --- | --- | --- | --- | --- | --- | --- | --- | --- | --- | --- | --- | --- | --- | --- | --- | --- |
| UM |  |  |  |  |  |  |  |  |  |  |  |  |  |  |  |  |  |  |
| 049H_H24-C.g515 | Field | na | NTS | DBLα | CIDRα | DBLβ | DBLβ | DBLy | DBLy | DBLδ | CIDRβ |  |  |  |  |  |  |  |
| QT0036-CW.g1117 | Field | na | NTS | DBLα | CIDRα | DBLβ | DBLβ | DBLy | DBLδ | CIDRβ |  |  |  |  |  |  |  |  |
| 024H_H08-C.g133 | Field | na | NTS | DBLα | CIDRα | DBLδ | CIDRβ | DBLz | DBLε |  |  |  |  |  |  |  |  |  |
| 044H_H24-C.g315 | Field | na | ATS |  |  |  |  |  |  |  |  |  |  |  |  |  |  |  |
| 065H_H16-C.g1122 | Field | na | NTS | DBLα | CIDRα | DBLβ | DBLδ | CIDRγ | DBLy | DBLz | ATS |  |  |  |  |  |  |  |
| CM |  |  |  |  |  |  |  |  |  |  |  |  |  |  |  |  |  |  |
| 074H_H08-C.g299 | Field | na | CIDRβ |  |  |  |  |  |  |  |  |  |  |  |  |  |  |  |
| PF07_0050 | PlasmoDB | na | NTS | DBLα0.18 | CIDRα6 | DBLβ6 | DBLy9 | ATS |  |  |  |  |  |  |  |  |  |  |
| SA |  |  |  |  |  |  |  |  |  |  |  |  |  |  |  |  |  |  |
| QT0009-CW4.g3425 | Field | na | NTS | DBLα | CIDRα | DBLδ | CIDRβ |  |  |  |  |  |  |  |  |  |  |  |
| 574984302 | NCBI | na | NTS | DBLα | CIDRα | DBLβ | DBLδ | CIDRγ |  |  |  |  |  |  |  |  |  |  |
| 069H_H00-C.g949 | Field | na | DBLβ | DBLβ | DBLδ |  |  |  |  |  |  |  |  |  |  |  |  |  |
| 069H_H00-C.g150 | Field | na | CIDRα | DBLβ3 | DBLy | DBLδ | CIDRγ |  |  |  |  |  |  |  |  |  |  |  |
| 061H_H16-C.g486 | Field | na | NTS | DBLα | CIDRα | DBLβ | DBLy | DBLδ | CIDRγ |  |  |  |  |  |  |  |  |  |
| QT0017-C.g13 | Field | na | NTS | DBLα | CIDRα | DBLβ | DBLy | DBLz | DBLε |  |  |  |  |  |  |  |  |  |
| 572SJ-C.g456 | Field | na | NTS | DBLα | CIDRα | DBLβ | DBLy | DBLy | DBLδ | CIDRβ |  |  |  |  |  |  |  |  |
| IT4var15 | VarDom | CD36 | NTS | DBLα0.8 | CIDRα3.5 | DBLβ3 | DBLδ1 | CIDRβ1 | ATS |  |  |  |  |  |  |  |  |  |
| 069H_H00-C.g45 | Field | na | DBLy | DBLδ | CIDRβ |  |  |  |  |  |  |  |  |  |  |  |  |  |
| 025H_H24-C.g225 | Field | na | ATS |  |  |  |  |  |  |  |  |  |  |  |  |  |  |  |
| QT0001-C.g391 | Field | na | NTS | DBLα | CIDRα | DBLβ | DBLδ | CIDRγ | DBLε | DBLz | DBLε | DBLβ | DBLδ | DBLε | DBLy | DBLz | DBLε | ATS |
| 069H_H00-C.g1030 | Field | na | NTS | DBLα | CIDRα | DBLβ | DBLy | DBLδ | CIDRβ | DBLβ | DBLy |  |  |  |  |  |  |  |
| QT0032-CM.g2637 | Field | na | NTS | DBLα | CIDRα | DBLβ | DBLδ | CIDRγ |  |  |  |  |  |  |  |  |  |  |
| 574984185 | NCBI | na | NTS | DBLα | CIDRα | DBLβ | DBLy | DBLz | DBLε | ATS |  |  |  |  |  |  |  |  |
| 610SJ-C.g1 | Field | na | ATS |  |  |  |  |  |  |  |  |  |  |  |  |  |  |  |
| DD2var29 | VarDom | CD36 | NTS | DBLα0.16 | CIDRα3.4 | DBLδ1 | CIDRβ1 | DBLy10 | ATS |  |  |  |  |  |  |  |  |  |
| 073H_H08-C.g576 | Field | na | NTS | DBLα | CIDRα | DBLy | DBLδ | CIDRβ | DBLε | DBLz | ATS |  |  |  |  |  |  |  |
| PFCLINvar21 | VarDom |  | DBLδ1 | CIDRβ1 | ATS |  |  |  |  |  |  |  |  |  |  |  |  |  |
| CM + SA |  |  |  |  |  |  |  |  |  |  |  |  |  |  |  |  |  |  |
| 069H_H08-C.g1025 | Field | na | DBLα | CIDRγ | DBLβ | DBLy | DBLy | DBLy | DBLy |  |  |  |  |  |  |  |  |  |
| DD2var28 | VarDom | ? | NTS | DBLα0.4 | CIDRα6 | DBLβ5 | DBLy18 | DBLε8 | ATS |  |  |  |  |  |  |  |  |  |
| 029H_H00-C.g187 | Field | na | NTS | DBLα | CIDRα | DBLδ | CIDRβ |  |  |  |  |  |  |  |  |  |  |  |
| 712SJ-C.g117 | Field | na | NTS | DBLα | CIDRα | DBLβ | DBLy | DBLz |  |  |  |  |  |  |  |  |  |  |
| 579318932 | NCBI | na | NTS | DBLα | CIDRα | DBLδ | CIDRβ | ATS |  |  |  |  |  |  |  |  |  |  |
| QT0033-CW3.g12 | Field | na | DBLβ | DBLβ | DBLδ | CIDRγ |  |  |  |  |  |  |  |  |  |  |  |  |

|  |  |  |  |  |  |  |  |  |  |  |
| --- | --- | --- | --- | --- | --- | --- | --- | --- | --- | --- |
| 069H_H00-C.g62 | Field | na | NTS | DBLα | CIDRα | | | | | |
| 021H_H00-C.g675 | Field | na | NTS | DBLα | CIDRδ | DBLβ | DBLβ | DBLγ | DBLδ | CIDRβ |

ICAM 1 binding + reference sequence

The PfEMP1 containing ICAM-1 binding pattern were associated to several protein groups, and matching with 5/9 of SA samples, 3/4 CM, and 4/9 UM samples (Figure 3). However, no peptide was identified corresponding directly to the pattern. We performed a 3D structure prediction (using 3D7 template available in PDB) from all DLBβ identified within the sequences obtained from WGS and identified in LC-MS/MS. Average quality score was 0.554 (± 0.047). RMSD calculation showed that the obtained DBLβ structure are highly similar (Figure 6 A).

**Figure 6:**
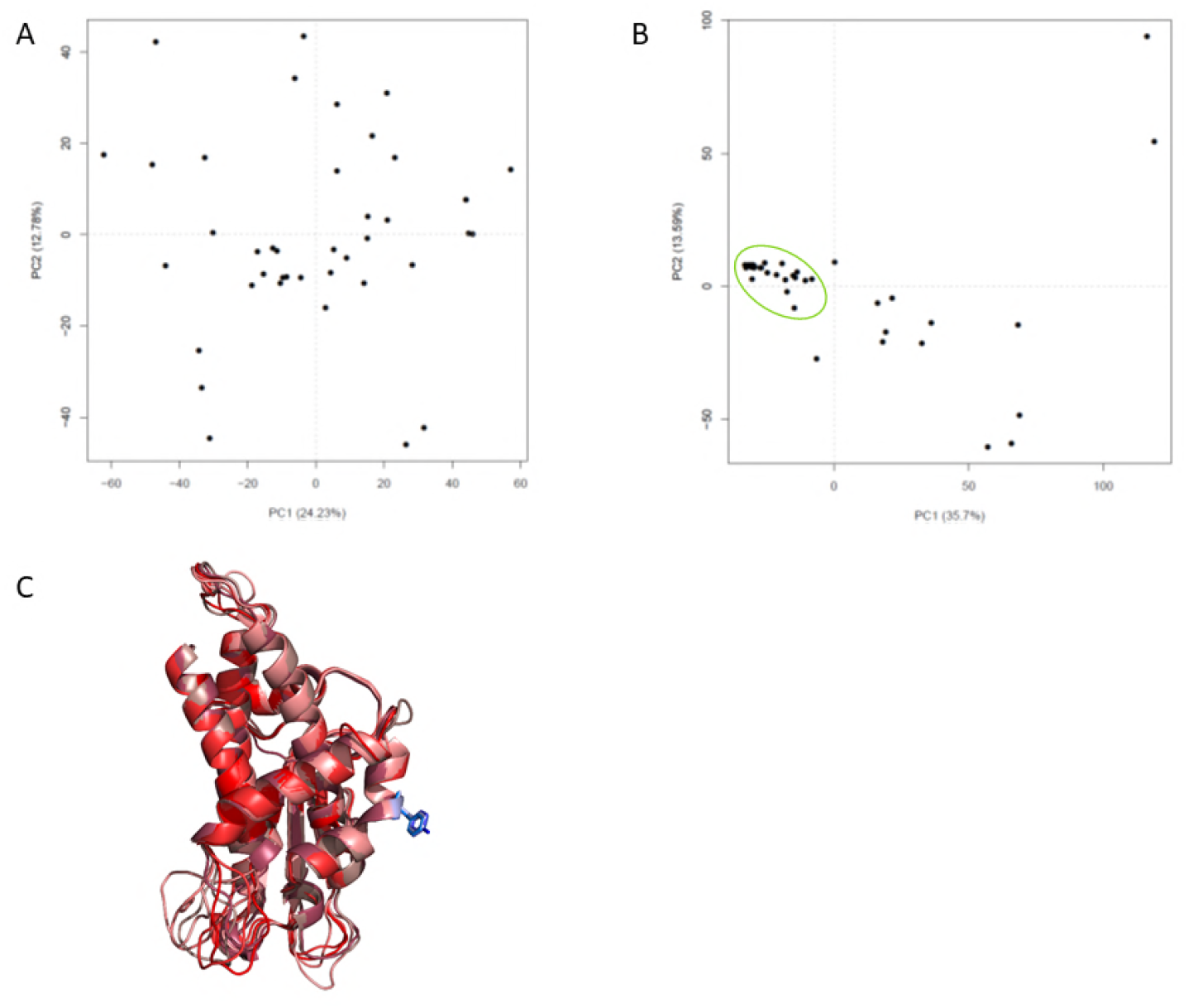
3D representation of the CIDRα associated to DBLβ containing ICAM-1 binding pattern. A) PCA representation of DBLβ RMSD values obtained after structure modelling. B) PCA representation of CIDRα RMSD values obtained after structure modelling. Green circle highlight sequences presenting CIDRα binding EPCR C) Juxtaposition of the structure predicted from the 6 CIDRα associated to DBLβ containing ICAM-1 binding pattern. Amino acid known for binding to EPCR is highlighted in blue. The two front α helix are folded accordingly to the structure from Hb3var3 and ITvar7, which bind EPCR. Structure have been drawn using Pymol.

Focusing on CIDRα, we were able to produce acceptable (quality score 0.470 ± 0.092) folding for all sequences except two which were excluded from the clustering. As expected, we observed that CIDRα1-like sequences and CIDRα “non α1” were clustered respectively (Figure 6 B). Using PyMol visualization of predicted 3D structure, we focused on the two α helix involved respectively in CD36 binding and interaction with EPCR (35). Among the CIDRα sequences, 14 presented a CIDRα “non α1” helix fold, among which 12 were folded correctly.

We then predicted the structure from CIDRα linked to DBLβ presenting ICAM-1 binding predicted pattern – 6 PfEMP1 proteins containing DBLβ-ICAM-1 binding pattern (Figure 6C). These CIDRα presented a phenylalanine (5/6 sequences) or tyrosine (1/6 sequences) in position 187 (1/6 sequences), 188 (3/6 sequences), 184 (1/6 sequences), 190 (1/6 sequences) from the corresponding CIDRα (Figure 6B). These amino acids were juxtaposed and exposed similarly to the phenylalanine involved in EPCR binding from Hb3var3 CIDRα 3D structure.

### Genomic and transcript data for specific protein identification

Since we had access to preserved RNA for 2 samples in addition to RNAsequencing data (CM04 and SA06 at 3 additional sampling time each: H0 (diagnosis of malaria) and post diagnosis sample, H16, H24 for CM04 and H0, H8, H16, for SA06), we selectively designed primers that would specifically amplify the PfEMP1 transcripts from the mass spectrometry matching sequences (Figure 1). Our goal was to assess which ones of these proteins are expressed, and their corresponding relative expression level using the Tu measurement method. Using alignment tools, we managed to find discriminant primers between the 6 identified sequences for sample SA06 (g45, g949, g1030, g1025, g150 and g62) and the 3 sequences (g276, g299 and g522) for sample CM04. Identified primers are presented in Supplementary data table 1 and quantification on Figure 7 for sample SA06. The amplification profile of reference genes was not satisfying for CM04-H0 and SA06–H08 samples, which were excluded from the analysis. For patient CM04, at H16, transcript expression was at the detection level threshold and no major transcript was identified. For patient CM04, at H24, the major transcript is g276 followed by g299 and g522 (Tu respectively: 43.26 – 4.24 – 1.32). For sequences that displayed different Melting Temperature (Tm), selective sequencing did not reveal differences between sequences, except for CM04 g276 sequence amplified with the designed primers.

**Figure 7:**
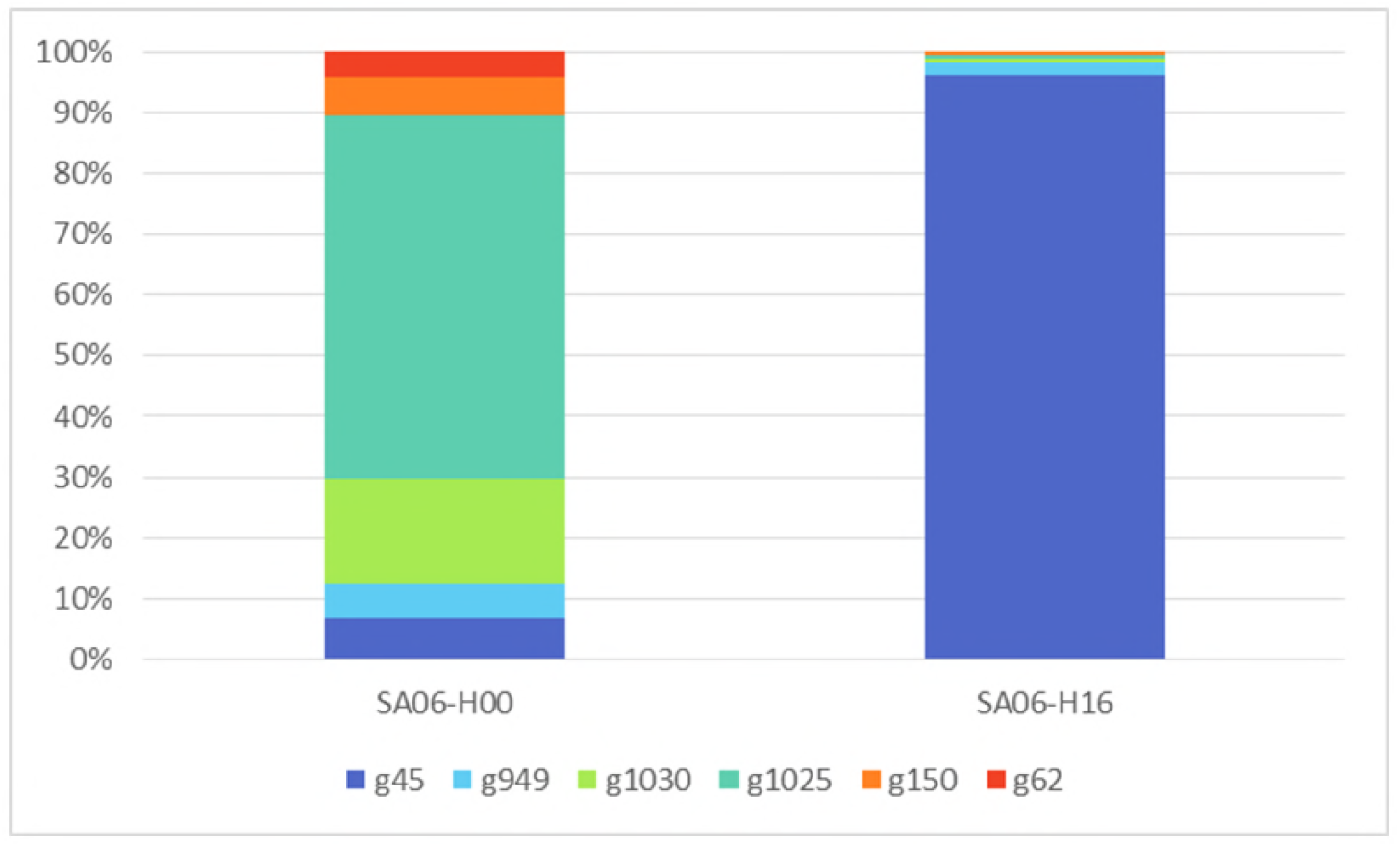
Transcript proportion for sample SA06, identified by selective RT-qPCR. Expression of *var* transcript within sample SA06 at diagnosis and 16h post treatment initiation. The relative percentage of each transcript is presented. Each transcript expression has been evaluated using dedicated primers specific to the newly identified *var* sequences. Using two reference genes for quantification, we identified the transcript g45 as the major expressed transcript for SA06 (at H16) samples, followed by transcript g949, g1030, g1025, g150, g62 (Tu respectively 3114.96 – 75.85 – 18.19 – 17.81 – 13.98 – 1.90). In the H0 sample, the following transcript were identified g1025, g1030, g45, g150, g949, g62 (Tu respectively: 23.59 – 6.77 – 2.64 – 2.41-2.25 – 1.66)

## Discussion

The evolution of *P. falciparum* infection from uncomplicated forms of the disease to cerebral malaria, the most fatal, is a complex phenomenon which entails host and parasite factors, in addition to socio-economics settings (36). There are strong evidence that the PfEMP1 proteins are involved in the disease progression since they allow the parasite to bind to host endothelium (14). It is believed that a distinct subset of PfEMP1 proteins is involved in severe malaria (23, 37), most likely by providing to the parasite the ability to sequester to a given receptor. However, PfEMP1 identification in natural infection remained challenging, due to the large size of PfEMP1 and their high sequences diversity. Recently, Jespersen et *al* (23) provided a new insight towards *var* genes sequences expression analysis in patient’s sample using transcript reconstruction after DBLα barcoding. They confirmed the preferential expression of CIDRα associated with EPCR binding in severe malaria patients.

We used a mass spectrometry–based proteomic approach to analyse the *P. falciparum* proteome in the context of severe malaria (SA and CM) compared to UM. We aimed to accurately identify, at the protein level, the PfEMP1 sequence variants associated with diseases severity. To this end, we initiated a “proteogenomic” study of field samples. In collaboration with the WT Sanger institute, we performed whole genome sequencing of the samples and had access to the assembled *var* gene sequences of the samples. Second, we performed RNAsequencing of these samples and analysed by LC-MS/MS the corresponding matured parasites. RNAsequencing reads were aligned for *P. falciparum* transcripts expression both on reference transcriptome and sequences issued from WGS of our samples. Reference database allowed the identification of *var* transcripts; however, all mapped reads were in semi conserved ATS coding region. Protein identification was performed by matching mass-spectrometry data against a custom database containing the translated PfEMP1 sequences (Figure 1). As anticipated, most of the identified PfEMP1 (considering all PfEMP1 from all protein group) came from the newly added sequences to the database (10/96 were known sequences), confirming the validity of our approach.

The technical improvement allowed us to identify more proteins than previously published studies (34, 38), with higher sequence coverage. We identified a set of 55 PfEMP1 associated in protein group. We investigated the structure of theses sequences and found that the two main domain organisations were NTS-DBLα-CIDRα-DBLβ and NTS-DBLα-CIDRα-DBLδ. The high proportion of NTS-DBLα-CIDRα-DBLβ among expressed PfEMP1 identified in our samples compared to genomic sequences within the same sample pool reflects the preferential expression of the PfEMP1 containing this pattern compared with the other PfEMP1 protein sequences in the studied context. The CIDRα-DBLβ tandem is associated with the potential “double binding” PfEMP1 (20, 24), targeting both ICAM-1 (through DBLβ (20)) and EPCR (through CIDRα (22)) human endothelial receptor. PfEMP1 binding pattern to ICAM-1 was not identified with specific peptides; however, proteins containing the binding pattern were identified in our samples (all malaria clinical presentation). We were able to predict domains structure for the CIDRα from PfEMP1 identified within our sequences pool. The proportion of “non α1” CIDR sequences identified is lower than the corresponding sequences in parasite genomes, (CIDRα2-6 represent 85% of PfEMP1 in parasite genomes) (37), enforcing the idea of preferential expression from PfEMP1 encoding EPCR binding CIDRα within malaria clinical presentation. In addition, all sequences presenting CIDRα-DBLβ- [ICAM-1 binder] harboured EPCR binding like CIDRα, enforced by 3D representation of domain structure.

For two isolates (SA09 and CM04), the identified PfEMP1 proteins were directly and unambiguously associated to transcript from the same isolate. To ensure whether several PfEMP1 proteins were expressed in the sample or if this was due to the identification method (peptides mapping sequences), we performed selective sequencing of the corresponding mRNA within the same sample. We managed to identify the most expressed transcript for one (CM04-H24), and the two main expressed transcripts for the other (SA09-H16). However, transcript corresponding to the other identified proteins existed at low expression level. It seems legitimate that along multiplication cycles, parasites express different PfEMP1. Furthermore, natural infection are often polyclonal and patients in endemic areas harbour usually 2 to 3 clones (39). However, we identified more sequences per sample than clones, strengthening the idea of simultaneous expression of several sets of PfEMP1 proteins by the parasite population within an individual. In addition, there are less PfEMP1 proteins identified in mass spectrometry compared to RNA sequencing/RT-qPCR approach. It seems plausible that, for technical reasons, PfEMP1 identification in LC-MS/MS provides a subset of the major proteins associated with each sample. In addition, all mRNA might not occur translation, as pseudogenes from *var* family are described. mRNA analysis reflects both coding and non-coding RNA expressed by a sample.

Focusing on patient’s group (CM, SA and UM), no consensus or highly similar sequence was obtained for a given clinical outcome of *P. falciparum* infection. Most sequences were shared between patient groups, since their identification relies mostly on reads mapping in the semi-conserved region ATS. We could not specifically identify sequences coding for the ICAM-1 binding pattern or the CIDRα involved in EPCR binding within severe malaria group. However, for sample SA06, one of the major proteins and transcript identified corresponds to a sequence expressing the binding pattern for ICAM-1.

We were not able to attribute domains subtype to the obtained sequences. The high diversity of PfEMP1 protein sequences compared with the repository database prevented us from assigning a subdomain to every domain. This limitation did not allow us to specifically identify subtypes of domains, in relation to reference sequences, associated with *P. falciparum* infection severity. However, selective alignment from obtained peptides to reference sequences repository allowed to identify domain subtypes for 36% (40/110) of identified peptides. This difficulty to attribute a domain may be the consequence of either: the peptide being shared between two or more domains, or the peptide has never been previously identified thus absent from the database (Figure 2). Variant protein identification in mass spectrometry is challenging, since protein identification is the reflection of the quality of the database available. Restrictive use of reference sequences from public repositories reduces the “chance” of protein identification, but we ride out this challenge with our home-custom sequence database obtained through whole genome sequencing of our samples. The variability of PfEMP1 primary sequence was overcome using protein structure modelling, which allowed us to classify the CIDRα identified within predicted binding target. We confirmed that PfEMP1 protein sequences which display ICAM-1 binding pattern present CIDRα domains with EPCR binding folding.

This study highlights the perspective offered by proteogenomic approaches for VSA studies. Using a home-made database containing PfEMP1 sequences obtained from WGS of the corresponding samples, we managed to identify specific PfEMP1 pattern associated with each sample, where RNAsequencing alone could not discriminate the expressed PfEMP1 within a sample. However, the sequence coverage of the identified proteins remained limited. Further attempts should focus on increasing the number of identified peptides per protein, using for example selective enrichment in membrane proteins. The identification of new PfEMP1 variants using whole genome sequencing and the confirmation of their expression at the protein level is, to our knowledge, the first report using field isolates.

In conclusion, we identified PfEMP1 proteins expressed by parasite in patients presenting several forms of malaria. This is the first report of PfEMP1 direct identification and is providing insight towards malaria pathogenesis understanding. PfEMP1 are adhesin which mediates iE adhesion to the host endothelium. The high proportion of CIDRα among the identified sequences enforce the idea that iE sequestration occurs either through CD36 binding, or EPCR binding, pending of clinical presentation (22, 40). We also preferentially identified PfEMP1 protein harbouring DBLβ, among which 20% (6/30 identified DBLβ) displayed the binding pattern for ICAM-1. This strengthen the hypothesis that DBLβ is involved in the disease development, as demonstrated with antibodies against DBLβ in Tanzania (41) and Papua New Guinea (42). Antibodies against full length DBLβ3_PF11_05_2_1_ are associated with reduction risk of severe malaria and protection against clinical infection (42). We demonstrated that predicted 3D structure from all DBLβ identified in the studied followed closely DBLβ3_PF11_0521_ structure, enforcing the idea of DBLβ3_PF11_0521_ as a vaccination target to prevent severe malaria. Further studies to identify immune response to DBLβ3PF11_0521, in combination with different CIDRα subtype are needed. There is little doubt that upcoming strategies to prevent severe malaria will target DBLβ3PF11_0521 in combination with EPCR binder CIDRα in a multi epitope vaccination strategy.

Our study opens opportunities to identify PfEMP1 variants and later implement these newly identified sequences in PfEMP1 based vaccine development strategies. However, our approach was restricted to one geographic area (Benin, West Africa) and included a limited number of patients. Further studies should include patients from various *P. falciparum* endemic areas to better represent PfEMP1 associated within *P. falciparum* disease in general and specifically to severe malaria.

## Material and Methods

### Ethic statement

Ethical clearance was obtained from the Institutional Ethics Committee of the faculty of health science at the Abomey-Calavi University in Benin (clearance n°90, 06/06/2016). Before inclusion, written informed consent was obtained from children’ guardians. Patients were treated in accordance to the national malaria program policy. The methods were carried out in accordance with the relevant guidelines and regulations.

### Sample collection

Patients under age 5, presenting severe malaria were included in the Lagune Mother and Child Hospital in Cotonou, Benin. Two distinct clinical groups were constituted, as following. CM was defined as a *Plasmodium falciparum* infection associated with a coma (Blantyre score ≤2) and the absence of meningitis detected by CSF count and culture. SA was defined as a *P. falciparum* infection associated with Hb < 5g/dL, measured using Hemocue^®^ device (Radiometer). UM patients were included in Saint-Joseph Hospital, in Sô-Ava, Benin (16 km from Cotonou). UM was defined as a *P. falciparum* infection with fever, in the absence of any other complication. The study was conducted at the rainy season (May – August) 2016.

Five mL of peripheral whole blood were collected on EDTA, (with additional sampling at 8h, 16h and 24h post treatment initiation for CM and SA patients). Parasite density was evaluated with Giemsa-stained thick blood smear. Only pure *P. falciparum* infection were retained for the study. Samples were depleted from white blood cells using a gradient based separation technique Ficoll (GE Healthcare Life Science).

### Whole genome sequencing

Fifty μL of erythrocyte’s pellet was extracted using DNEasy Blood kit (Qiagen). WGS was performed by the Malaria Genomic Epidemiology Network (MalariaGEN) at the Welcome Trust Sanger Institute (Hinxton, UK).

### Transcriptome studies

Ring staged parasite were preserved in 5 volumes of pre-warmed (37°C) TriZol (Life Technology), vortexed then immediately frozen at −80°C until further utilization. RNA was extracted as described (43), then digested with DNAse I (Qiagen) and purified using RNEasy MinElute Cleanup kit column (Qiagen). Absence of genomic DNA was assessed by the absence of amplification of p90 gene in qPCR from the direct use of the RNA extract, using Syber Green reagent (Life Technology) on Rotorgene 6000 (Corbett Technology). RNA quality was assessed using PicoChip Agilent 2100TM Bioanalyzeur profile (Agilent). Only RNA presenting a RNA Integrity Number (RIN) > 7 were retained for downstream analysis (44). Sequencing technology used was an Illumina NexSeq500 (IPS2 POPS platform). RNA-seq libraries were performed by TruSeq Stranded mRNA protocol (Illumina^®^, California, U.S.A.). The RNA-seq samples have been sequenced in paired-end (PE) with a sizing of 260 base pairs and a read length of 150 bases. 54 samples by lane of NextSeq500 using individual bar-coded adapters and giving approximately 5 million of PE reads by sample are generated.

### RNA-sequencing data bioinformatics treatment and analysis

RNA-Seq preprocessing includes trimming library adapters and performing quality controls. The raw data (fastq) were trimmed with Trimmomatic (45) tool for Phred Quality Score Qscore >20, read length >30 bases, and ribosome sequences were removed with tool sortMeRNA (46). Reads were mapped against the *Plasmodium falciparum* 3D7 reference genome (PlasmoDB v35) and against the reconstructed sequences from WGS, using STAR (version 2.5.4b) with default parameters. A sequence was considered as successfully sequenced if more than 70% of the transcript was covered with at least 5 reads.

### Obtention of mature forms from *P. falciparum*

Blood samples were matured *in vitro* for 18 to 32 hours in RPMI medium supplemented with human serum and Albumax (Gibco) and preserved after MACS™ (Myltenyi Biotech) enrichment as described (33).

### Protein extraction and sample preparation for mass spectrometry analysis

Whole cell infected erythrocyte lysates were solubilized and digested in solution using trypsin (Promega, sequencing Grade). Briefly, 50 μg of proteins from whole cell lysates were diluted to 25 μl in solubilization buffer (1% sodium desoxycholate, 100 mM Tris/HCl, pH8.5, 10mM TCEP, 40 mM chloroacetamide), heated for 5 min at 95°C and sonicated three times for 30 s. Once at room temperature, extracts were diluted with 25 μl Tris-ACN buffer (50mM Tris/HCl pH 8.5, 10%ACN). Collected peptides were fractionated in 5 fractions per sample by strong cationic exchange (SCX) StageTips (47).

### Mass spectrometry analysis

Mass Spectrometry (MS) analysis were performed on a Dionex U3000 RSLC nano-LC-system coupled to an Orbitrap-fusion mass spectrometer (Thermo Fisher Scientific) as described (48). Peptides were separated on a C18 reverse-phase resin (75-μm inner diameter and 15-cm length) with a 3-hr gradient. The mass spectrometer acquired data throughout the elution process and operated in a data-dependent scheme.

### Accession number

The raw reads were available under the accession number listed in supplemental data 5. VarDOM server for *var* genes is available at the following address www.cbs.dtu.dk/services/VarDom/. PlasmoDB reference sequences are available on EupathDB portal dedicated to *Plasmodium* (http://plasmodb.org/plasmo/).

### Database construction

To get an accurate identification of proteins, a database was created as follows: Since our samples have been sequenced by the MalariaGen consortium (UK), whole genome sequences were available. Reconstructed *var* genes were kindly provided by Thomas Otto, Matt Berriman and Chris Newbold from the Welcome Trust Sanger Institute. We concatenated *P. falciparum* proteins sequences from PlasmoDB (v35) Uniprot and NCBI. We added our own PfEMP1 sequences, obtained after *in silico* translation from *var* genes reconstruction. Duplicate sequences were removed. FASTA sequence headers were uniformed to facilitate MaxQuant sequences analysis.

### Raw data processing

The mass spectrometry data were analyzed using Maxquant version 1.5.2.8 (49). The database used was our homemade database (see above section) and the list of contaminant sequences from Maxquant. The precursor mass tolerance was set to 4.5 ppm and the fragment mass tolerance to 0.5 Da. Carbamidomethylation of cysteins was set as constant modification and acetylation of protein N-terminus and oxidation of methionine were set as variable modifications. Second peptide search was allowed, and minimal length of peptides was set at 7 amino acids. False discovery rate (FDR) was kept below 1% on both peptides and proteins. Label-free protein quantification (LFQ) was done using both unique and razor peptides. At least 2 such peptides were required for LFQ. The “match between runs” (MBR) option was allowed with a match time window of 1 min and an alignment time window of 30 min.

For analysis, LFQ results from MaxQuant were imported into the Perseus software (version 1.5.1.6). Reverse and contaminant proteins were excluded. Only proteins from *P. falciparum* were selected for further analysis. We then focused on the membrane associated and putative proteins from *P. falciparum*.

### Protein sequence analysis and structure prediction

PfEMP1 sequences were aligned using the VarDom server (available at http://www.cbs.dtu.dk/services/VarDom/) for domain type identification (11). We specifically searched the pattern for ICAM-1 binding I[V/L]x_9_N[E]GG[P/A]xYx_27_GPPx_3_H (20) in the identified PfEMP1 sequences using the ProSite online interface (50).

Using domain limitation performed with VarDOM server, we predicted the structure of the CIDRα and DBLβ domains identified within our sequences. Structure were generated using Modeller (v9.20.) (51). We selected templates from the Protein Data Bank (52) using BLAST algorithm. Template selection was performed manually using BLAST results, sequence coverage within each template and template quality. DBLβ structure prediction was performed using PDB 5mza (PF11_0521). For the CIDRα domain, the following templates were used: 4v3d (from HB3var03), 4v3e (from IT4var07), 3c64 (MC179) and 5lgd (from MCvar1). Template and query quality scores were calculated using ProQ3 online tool. For 5mza the quality score was 0.560. For 4v3d, 4v3e, 3c64 and 5lgd, quality scores were respectively 0.484, 0.585, 0.396 and 0.540. For each domain, 5 models have been generated, and 4 additional models were generated after loop refining. Template and query mapping has been performed using align2D tool from Modeller script. For structure modelling using several templates, we used salign script (part of Modeller script).

The choice of the best model was made based on the Discrete Optimized Protein Energy (DOPE) score values. In order to analyze and compare the modeled structures were compared using the Bio3D v 3.2 R package (53). Structure comparison was based on Root-Mean-Square Deviation (RMSD) calculated using Cα distance from predicted structures. Obtained RMSD values were clustered using Bio3D R package (53).

### Statistical analysis

Patient’s samples information’s were compared between the 3 patient’s groups (UM, CM and SA) using one-way ANOVA. Bonferroni’s Multiple Comparison Test was applied for individual group comparison. We considered a *p* value < 0.05 as significant. Qualitative data were compared with Chi Squared test using contingency table. All analyses were performed using Prism v5 (Graphpad).

### Primer design

We tried to specifically assign PfEMP1 expression within the samples. In addition to RNAsequencing data, we disposed of preserved RNA from 2 samples. Our strategy was as follows: first, we compared the sequences of the identified proteins using Multalin (54). Second, we designed specific primers targeting the sequences using Primer3Plus web interface (55). Finally, we assessed primers specificity using BLAST tools. (Primers are listed in Supplementary data 2).

### RT-qPCR and PCR

RNA was preserved using Trizol reagents (Life Technology) and RNA extraction was performed as described (43). cDNA was obtained using Superscript II (Life Technology) retrotranscriptase following the manufacturer’s instructions.

qPCR was performed using Power SYBR Green PCR Master Mix (Applied Biosystems) on Rotorgen (Corbett technology) as described (34). Melting stage was set up as default on the Rotorgen software. PCR specificity was assessed by the melting temperature (Tm). We considered that a sequence is unique for Tm included in the range Tm +/− 0.5°C. Transcript expression was quantified using the Transcript Unit (Tu) method, as described by Lavsten et *al* (31). GAPDH and Seryl-tRNA synthetase transcripts were used as reference genes.

For transcripts which displayed a Tm difference upper than 0.5°C for a same primer set, we performed a conventional PCR using AmpliTaGold DNA polymerase (Applied Biosystem) with the following conditions: initial denaturation at 95°C for 10 min, followed by 52°C hybridization for 30s – 68°C elongation for 15 s, repeated 35 times. PCR products length was assessed using 2% agarose gel containing Syber Safe (Life Technology) at 1/10,000.

Amplicons were purified using the QIAquick PCR Purification Kit (Qiagen). The purified fragments were sequenced with BigDye Terminator v3.1 Cycle Sequencing Kit (Applied Biosystems) using g949, g150 (for patient SA06), and g299 primers (for patient CM04) (Supplementary data 1). The sequence reaction products were purified using the BigDye XTerminator^®^ Purification Kit (Applied Biosystems), in accordance with the manufacturer’s instructions. The purified products were sequenced using an ABI Prism 3100 analyzer (Applied Biosystems), and the sequences were analyzed using DNA Baser v4 (Heracle BioSoft).

## Acknowledgments

The authors would like to thank patients who participate in the study, the clinicians and nurses who were involved in patient’s inclusion. We especially acknowledge Dr Nadine Fievet help and counselling during the field study. We would like to thank Matt Berriman, Chris Newbold and Thomas Otto from the Welcome Trust Sanger Institute for the access to the unpublished PfEMP1 sequences. We also would like to thank Emilie-Fleur Gautier for her helpful advices in mass spectrometry, and Evangeline Bennana for her technical support in sample preparation. The authors thank Romain Coppée for his critical comments on the manuscript, and Antoine Claessens for his help in WGS. We also thank Feder for the opportunity to use Orbitrap Fusion mass spectrometer.

## Author’s contribution

CK, PD, FG and GB designed the study. CK, SE, JR, JMA and GB conducted the field study. CK, GB, CB, SG, CP and SH performed the experiments. CK, DR and EG analyzed the data. CK, GB and PD wrote the paper.

## Funding

This work was funded by Merieux Research Grant (awarded to Dr Gwladys Bertin for CIVIC project), by the Laboratoire d’Excellence GR-Ex, Paris, France, reference ANR-11-LABX-0051, that is funded by the program Investissements d’avenir of the French National Research Agency, reference ANR-11-IDEX-0005-02 and by NeuroCM project, that is funded by ANR-17-CE17-0001

The POPS platform benefits from the support of the LabEx Saclay Plant Sciences-SPS (ANR-10-LABX-0040-SPS).

CK is awarded a PhD Scholarship from French Minister of Research through MTCI Doctoral School (ED 563, Paris Descartes University).

## Supplemental material legends

### Supplemental material 1

Control transcript evaluation for RNAsequencing

### Supplemental material 2

Primers used for selective PfEMP1 sequencing

### Supplemental material 3

Identification of PfEMP1

### Supplemental material 4

Selective sequencing of *var* genes within field samples

### Supplemental material 5

ENA accession numbers from genomic datafiles

